# Multimodal Bonds Reconstruction Towards Generative Molecular Design

**DOI:** 10.1101/2025.05.06.652517

**Authors:** Jian Wang, Nikolay V. Dokholyan

**Affiliations:** Department of Neuroscience and Experimental Therapeutics, Penn State College of Medicine, Hershey, PA, 17033-0850, USA; Department of Biomedical Engineering, Pennsylvania State University, University Park, PA; Department of Chemistry, Pennsylvania State University, University Park, PA

## Abstract

Generative models such as diffusion-based approaches have transformed *de novo* drug design by enabling rapid generation of novel molecular structures in both 2D and 3D formats. However, accurate reconstruction of chemical bonds, especially from distorted geometries produced by generative models, remains a critical challenge. Here, we present YuelBond, a multimodal graph neural network framework for robust bonds reconstruction across three key scenarios: (i) recovery of bonds from accurate 3D atomic coordinates, (ii) reconstruction of chemically valid bonds in crude *de novo* generated compounds (CDGs) with perturbed geometries, and (iii) reassignment of bond orders in 2D topological graphs. YuelBond outperforms traditional rule-based methods such as RDKit, achieving 98.4% F1-score on standard 3D structures and maintaining strong performance (92.7% F1-score) on distorted CDGs, even when RDKit fails on most cases. Our results demonstrate that YuelBond enables accurate and reliable bond reconstruction from imperfect molecular data, bridging a critical gap in generative drug discovery pipelines.

## INTRODUCTION

In recent years, generative artificial intelligence—particularly generative adversarial networks^1^ (GANs) and diffusion models^2^—has revolutionized *de novo* drug discovery^3^ by enabling the rapid generation of massive novel molecular structures. These generative approaches are broadly categorized into two-dimensional (2D) molecular topology generation and three-dimensional (3D) molecular conformation generation. 2D molecular topology generation focuses on constructing molecular graphs or string-based representations (e.g., SMILES^4^/SELFIES^5^), while 3D molecular conformation generation aims to predict atomic coordinates in 3D space. For 2D generation, widely used methods include autoregressive models such as MolRNN^6^, BIMODAL^7^, and MolecularRNN^8^, variational autoencoders such as junction tree variational autoencoder^9^, GraphVAE^10^, and CGVAE^11^, GAN-based frameworks like LatentGAN^12^, druGAN^13^, and MOLGAN^14^, and flow-based architectures like GraphNVP^15^ and MoFlow^16^. Meanwhile, 3D generation primarily relies on diffusion models—such as DiffSBDD^17^ and DiffBP^18^—which often employ equivariant graph neural networks^19^ (EGNNs) to model molecular structures within protein binding pockets.

Despite their promise, these generative methods face critical limitations, particularly concerning chemical bond order accuracy. In 2D generation, predicted bond orders may be chemically invalid, while 3D generation typically outputs only atomic coordinates without explicit bond annotations. This workflow leads to two major challenges: (1) validation difficulties, as incorrect bond orders hinder reliable assessment of molecular validity^20,21^, and (2) degraded performance in downstream tasks, including molecular dynamics simulations^22,23^, docking studies^24,25^ and protein-small molecule binding prediction^26–28^, where bond order errors propagate into inaccurate force field parameterization and binding affinity predictions. Addressing these issues is crucial for bridging the gap between *in silico* molecule generation and real-world drug development applications.

Typically, bond orders are determined by hybridization analysis using bond lengths and angles^29^ or by comparison of atomic pairs with a known database^30,31^. They can be also assigned using chemical or length rules^32^. RDKit^33^, the most popular cheminformatics toolkit, uses the xyz2mol^34^ program to predict the bond order of the molecule from the 3D coordinates. xyz2mol determines bond orders from 3D coordinates through a rule-based approach. First, it identifies connected atoms using covalent radii. Then it calculates possible valence states and systematically tests bond order distributions between unsaturated atoms while respecting valence rules. The solution uses either combinatorial pairing or graph theory to find chemically valid configurations, optionally employing Hückel theory for ambiguous cases.

However, existing methods are not robust to geometric distortions in generated molecules. When the geometry is distorted, i.e., bond lengths and angles deviate from ideal values, even determining atomic connectivity becomes challenging. Predicting bond orders under such conditions is even more difficult. This problem was less critical before generative models were widely used in drug discovery, but as these models become increasingly prevalent, the issue is becoming more serious. On one hand, generative models must produce increasingly accurate 3D molecular structures. On the other hand, we must develop more robust approaches to infer molecular connectivity and bond orders directly from distorted 3D coordinates.

To address these limitations, we introduce YuelBond, a graph neural network (GNN)-based framework that infers bond orders from molecular representations, whether they are accurate 3D coordinates, generated noisy structures, or even mere 2D topological graphs. Unlike conventional GNNs that focus on node-level tasks, YuelBond is designed for edge-centric learning, explicitly modeling the bond between pairs of atoms using their interatomic distances, local atomic environments, and iterative message passing. This allows the model to learn subtle patterns associated with different bond types and generalize beyond handcrafted rules. YuelBond is evaluated across three increasingly challenging scenarios: (1) reconstruction of bond orders from accurate 3D atomic coordinates; (2) prediction from noisy 3D structures simulating conformations generated by generative models; and (3) reassignment of bond orders given only 2D molecular connectivity. Across these tasks, YuelBond demonstrates robust and accurate performance, surpassing traditional methods, especially under structurally noisy conditions where existing tools often fail. Furthermore, by outputting class probabilities, YuelBond provides a nuanced view of prediction uncertainty, which is an essential feature for downstream workflows that must reason over ambiguous or probabilistic bonding assignments. Taken together, our results show that YuelBond is a flexible and reliable tool for bond order inference across multiple molecular representations.

## RESULTS

### Bonds Reconstruction from Accurate 3D Coordinates

YuelBond is developed based on GNN, in which each molecule is represented as a graph with atoms serving as nodes. For every atom, we calculate its distances to all other atoms in the molecule, and any pair of atoms within 3 Å is connected by an edge (Figure 1). These pairwise distances are assigned as edge features. While conventional GNNs are typically used for node-level predictions^35–38^, we adapt the architecture to focus on predicting edge-level features, specifically the bond orders between atoms. To accomplish this, we modify the aggregation mechanism for edge updates (Methods). Instead of relying solely on the concatenation of features from the connected nodes, we additionally incorporate the distance between atoms and the previous edge features into the update function. This design allows the model to capture the local chemical environment around each bond more effectively. With each layer of message passing, the edge features are refined, thereby expanding the receptive field. At the final stage, a linear layer maps the updated edge features to one of four bond order categories: single, double, aromatic, or triple (Figure 2).

**Figure 1.**
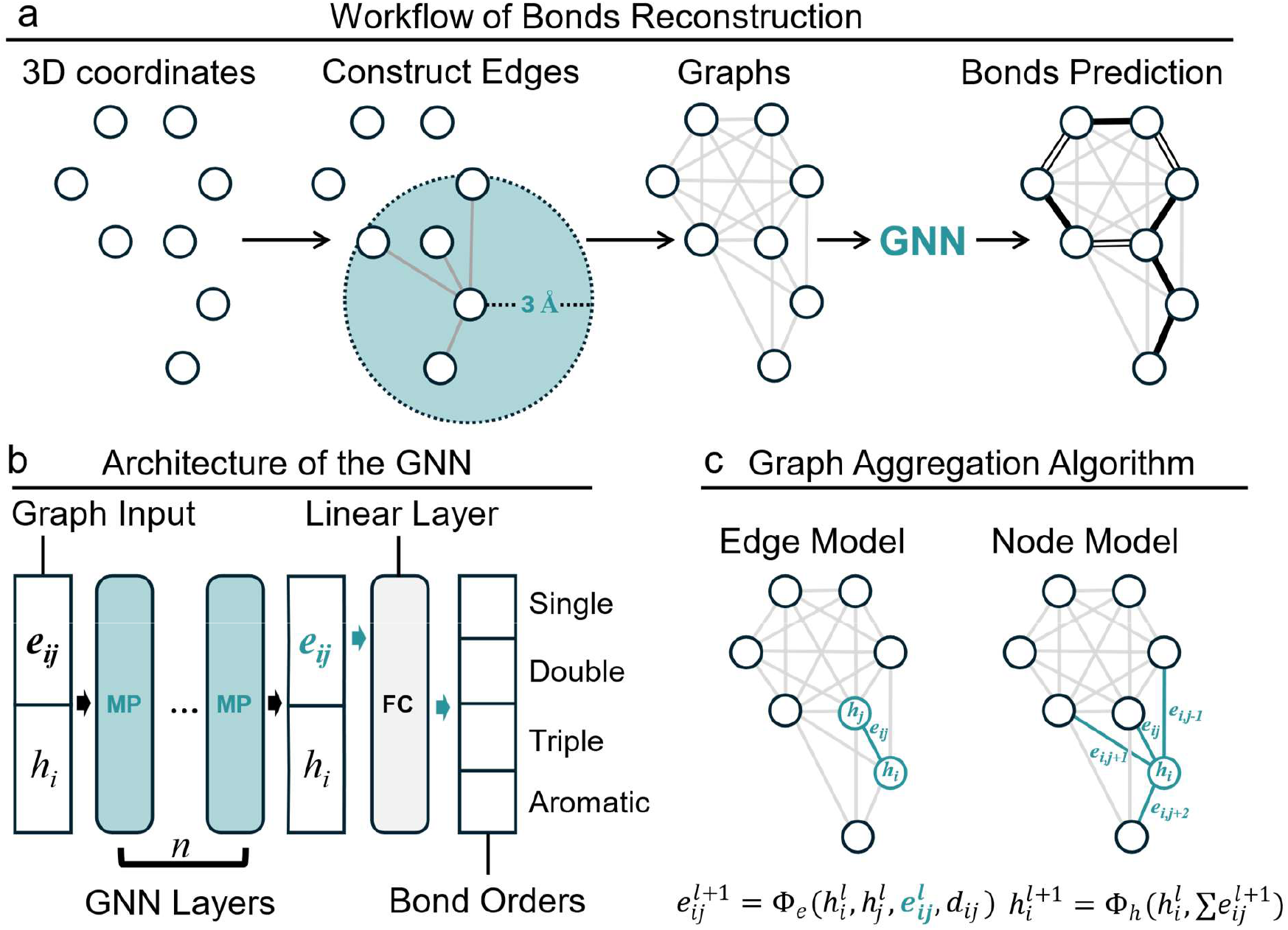
Workflow and architecture for bonds reconstruction using a graph neural network. (a) The process begins with 3D atomic coordinates as input, followed by edge construction to form a molecular graph, where edges are defined between atoms within a 3 Å cutoff distance. The graph, containing node (atom) and edge (interatomic) features, is then processed by a GNN to predict bond orders (single, double, triple, or aromatic). (b) The GNN architecture consists of graph input, linear layers, and multiple message-passing (MP) layers that iteratively update edge embeddings (*e*_*ij*_) and node states (*h*_*i*_) through message passing. (c) The edge features and node features are aggregated using the edge model and node model.

**Figure 2.**
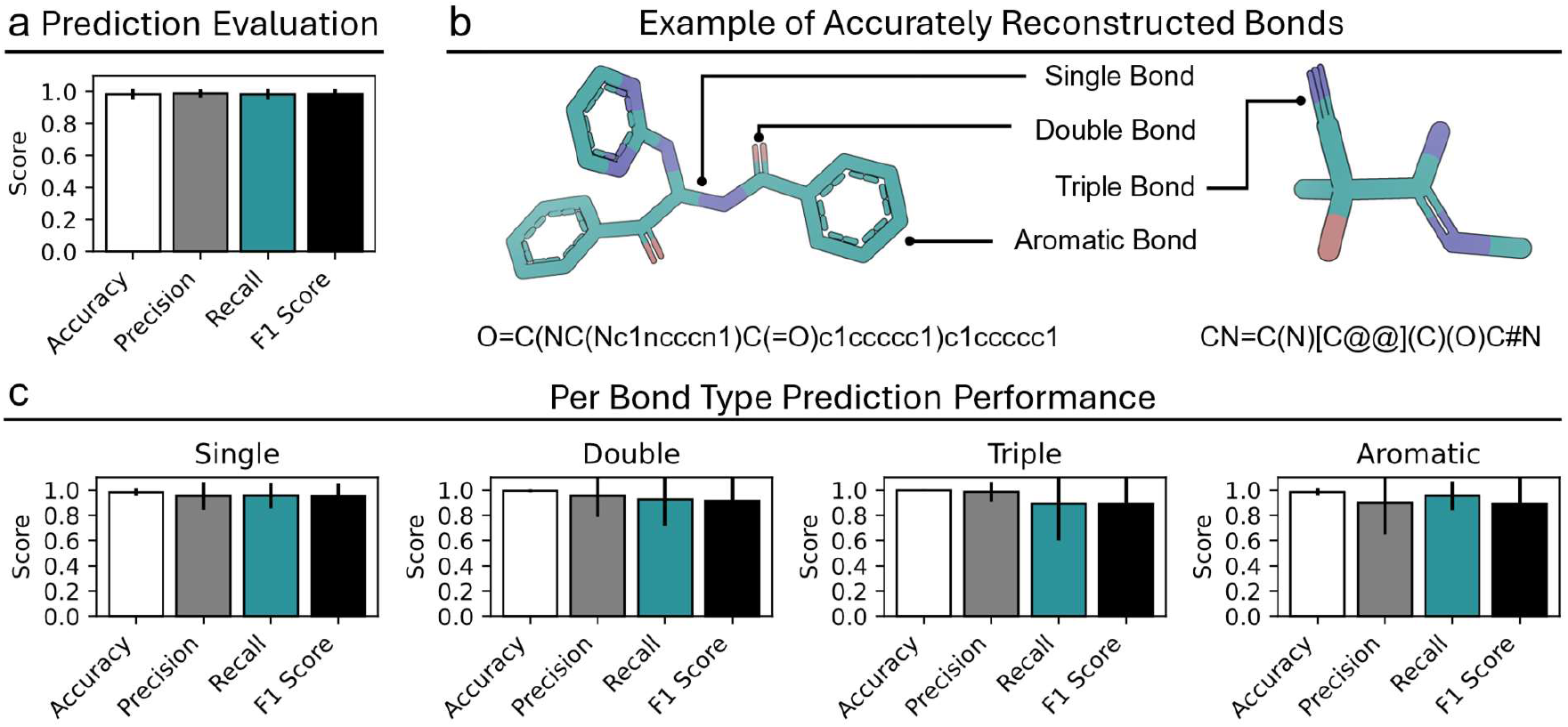
Performance Evaluation of Bond Order Prediction in Molecular Structures. (a) The average accuracy, precision, recall, and F1 score. (b) Examples of accurately predicted bond orders. (c) Per bond type prediction performance.

We trained and evaluated YuelBond using the Geometric Ensemble of Molecules (GEOM)^39^ dataset, which includes over 450,000 molecular structures. The dataset was randomly split into training, validation, and test sets. The validation set was used for tuning hyperparameters during training. To evaluate bond reconstruction performance, we first removed all bond annotations from the molecules and used YuelBond to reconstruct the bonds based on their 3D atomic coordinates. We then calculated accuracy, precision, recall, and F1-score on the test set. The model achieved an average accuracy of 98.2%, precision of 98.8%, recall of 98.2%, and F1-score of 98.4%, demonstrating a strong capability in recovering molecular bonding information (Figure 2a; Table S1).

We also analyzed YuelBond’s prediction performance across different bond types (Figure 2c; Table S1). For single bonds, the model achieved an average accuracy of 98.4%, precision of 95.4%, recall of 95.6%, and F1-score of 95.0%. For double bonds, the corresponding metrics were 99.6% accuracy, 95.4% precision, 92.6% recall, and 91.4% F1-score. Triple bond prediction yielded an accuracy of 99.94%, precision of 98.75%, recall of 89.4%, and F1-score of 89.4%. Aromatic bond prediction achieved 98.6% accuracy, 90.1% precision, 95.5% recall, and 89.4% F1-score. These results confirm that YuelBond is capable of accurately reconstructing diverse bond types from 3D atomic arrangements, including those that are challenging for traditional rule-based approaches.

The high performance on single and double bonds suggests that the model effectively learns typical interatomic distances and local environments associated with common covalent bonding. The slightly lower recall for triple bonds, despite high precision, indicates that the model tends to be conservative in assigning triple bonds, possibly due to their rarity and the strict geometric constraints required for their identification. This conservative behavior is preferable as misclassifying a non-triple bond as triple could result in larger errors in downstream tasks.

### Bonds Reconstruction from Crude *De novo* Generated Compounds

The goal of this work is to develop a model capable of reconstructing chemical bonds in crude *de novo* generated compounds (CDG), which often exhibit noisy or imprecise 3D structures. To simulate this scenario, we introduce controlled noise into the molecular coordinates of the GEOM dataset (Figure 3a; Methods), mimicking the structural distortion commonly found in generated molecules. We then employ our model to reconstruct bonds from these perturbed coordinates. Evaluation on the test set shows that the model maintains strong performance under noisy conditions, achieving average accuracy, precision, recall, and F1-score of 92.7%, 93.6%, 92.7%, and 92.7%, respectively (Figure 3b; Table S2).

**Figure 3.**
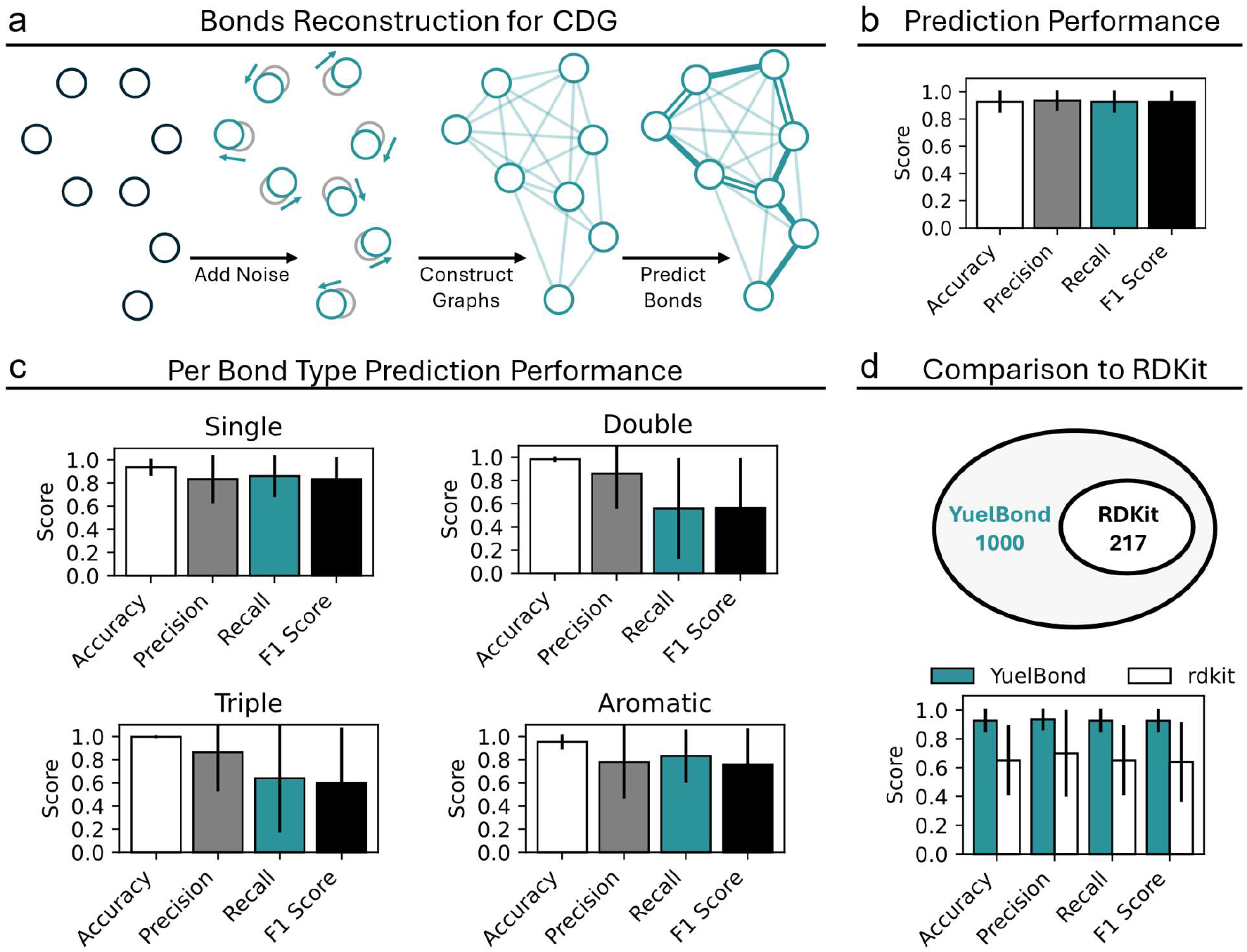
Performance Evaluation of YuelBond on the CDG Bonds Reconstruction Task. (a) Noise is introduced to the atomic coordinates in the compound, and edges are constructed from the perturbed atoms. Bonds are then reconstructed from these distorted compounds. (b) Evaluation of YuelBond’s performance in reconstructing the CDG bonds. (c) Performance of YuelBond for predicting each bond type. (d) Comparison of YuelBond’s performance with RDKit. YuelBond successfully processes all 1000 compounds in the test set, while RDKit can only process 217 compounds due to failures caused by distorted bond lengths and angles. YuelBond achieves approximately 0.8 in accuracy, precision, recall, and F1 score across all 1000 compounds, while RDKit obtains ~0.6 for these metrics on the 217 compounds.

We further analyzed the performance of YuelBond across different bond types (Figure 3c; Table S2). For single bonds, the model achieved an accuracy of 93.5%, precision of 83.4%, recall of 86.0%, and F1-score of 83.3%. Double bond prediction yielded 98.3% accuracy, 85.9% precision, 55.9% recall, and 56.3% F1-score. For triple bonds, the model achieved 99.8% accuracy, 86.5% precision, 63.9% recall, and 59.9% F1-score. Aromatic bond prediction produced 95.2% accuracy, 78.0% precision, 83.2% recall, and 75.5% F1-score.

These results show that the model retains high predictive accuracy for all bond types, but precision and recall vary with bond complexity. Single bonds are predicted most reliably, due to their high frequency and distinctive local geometry, even after perturbation. Aromatic bonds, often embedded in rings with recognizable topological features, are also well predicted despite coordinates noise. In contrast, double and triple bonds present greater challenges under distortion. Their prediction relies heavily on precise bond lengths and angles, which are features that become less informative when the structure is noisy. Consequently, the recall and F1-scores for these bond types are lower, reflecting increased false negatives.

To benchmark our model, we compared its performance with RDKit (Figure 3d; Table S2). Out of 1,000 test molecules with noisy coordinates, RDKit was able to process only 217, while it failed on the remaining 783 molecules. This limitation arises because RDKit relies on chemically valid input geometries and rule-based heuristics that become unreliable when the structure is distorted. Even among the 217 successfully processed molecules, RDKit achieved only 65.3% accuracy, 70.1% precision, 65.3% recall, and 64.2% F1-score – substantially lower than YuelBond. These results highlight the robustness of YuelBond in reconstructing bond orders from imprecise 3D structures, where traditional rule-based methods often fail.

### Bond Order Reassignment for 2D Generated Compounds

In the third scenario, we consider cases where generated molecules are represented solely as 2D graphs—that is, with atom connectivity but without 3D coordinate information (Figure 4a). This format is common for generative models that produce SMILES strings or molecular graphs, where the presence of a bond is known but the bond order is either missing or ambiguous. In such cases, reassignment of bond orders is a key step toward constructing chemically valid structures.

**Figure 4.**
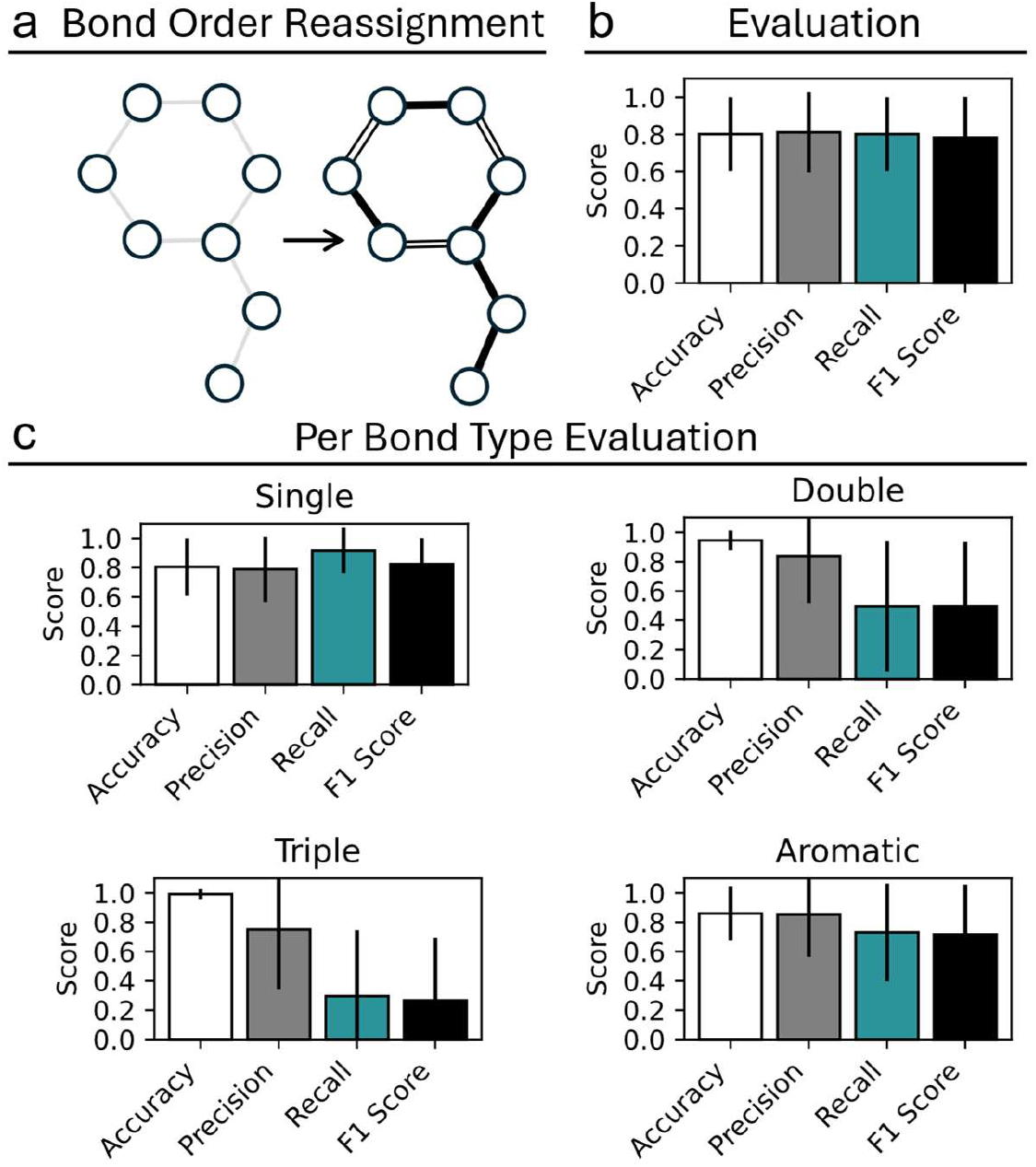
Performance Evaluation of YuelBond on the Bond Order Reassignment Task. (a) The bond connectivity is preserved, while the bond orders are reassigned across all bonds. (b) Evaluation of YuelBond’s performance in reassigning bond orders. (c) Performance of YuelBond for predicting each bond type.

To address this challenge, we applied YuelBond to infer bond orders based only on the atomic connectivity. When evaluated on the test set, the model achieved average accuracy, precision, recall, and F1-score of 80.1%, 81.1%, 80.1%, and 78.3%, respectively (Figure 4b; Table S3). Compared to the previous scenarios that incorporated 3D information, these values are lower, reflecting the increased complexity of bond order prediction without geometric features such as bond lengths and angles.

We further examined the model’s behavior across different bond types (Figure 4c; Table S3). For single bonds, YuelBond achieved high recall (91.7%) and a solid F1-score (82.4%), reflecting the consistency and prevalence of these bonds in molecular graphs. For double and triple bonds, accuracy remained high (94.6% and 99.2%, respectively), but recall was lower due to their relatively low frequency. This is expected, as higher-order bonds often exhibit subtler patterns that are more challenging to distinguish based on connectivity alone. Notably, the high accuracy for triple bonds indicates strong capability in avoiding false positives, while the lower recall highlights a more conservative prediction strategy that favors precision over overprediction. The lower recall for double and triple bonds, relative to single and aromatic bonds, is largely attributable to their rarity in the dataset. Since the overall number of negative cases (e.g., bonds that are not triple) is much larger, even a conservative predictor can yield high accuracy. Aromatic bonds were predicted with balanced performance, achieving 86.0% accuracy and an F1-score of 71.6%, suggesting that their unique connectivity patterns are reasonably well captured by the model.

Molecules with identical 2D connectivity may correspond to different chemically valid bond order assignments due to variations in local electronic environments. A simple example is the distinction between pyridine and its N-oxide form, where the connectivity remains unchanged but the bond orders around the nitrogen atom differ. Similarly, consider the case of a carbon-carbon bond, where a single bond, a double bond, and a triple bond could all appear as valid assignments depending on the electronic environment and the molecule’s resonance structure. In compounds like alkynes or alkenes, the bonding pattern can shift between single, double, or triple bonds, even if the connectivity graph appears identical in 2D representation. These subtle differences are inherently challenging to resolve based on connectivity alone. Nonetheless, YuelBond demonstrates the ability to make chemically meaningful and robust predictions for bond order reassignment.

### Bond Order Probability Prediction

While the overall prediction accuracy for double and triple bonds is lower in the bond order reassignment scenario (Figure 5a&b), it is informative to examine the confidence in the predictions. Since YuelBond outputs raw logits for each bond order class, we apply the *softmax* function to convert them into probabilities to reflect the confidence in assigning each possible bond order. These probabilities are meaningful as multiple bond types may appear plausible based on 2D connectivity alone.

**Figure 5.**
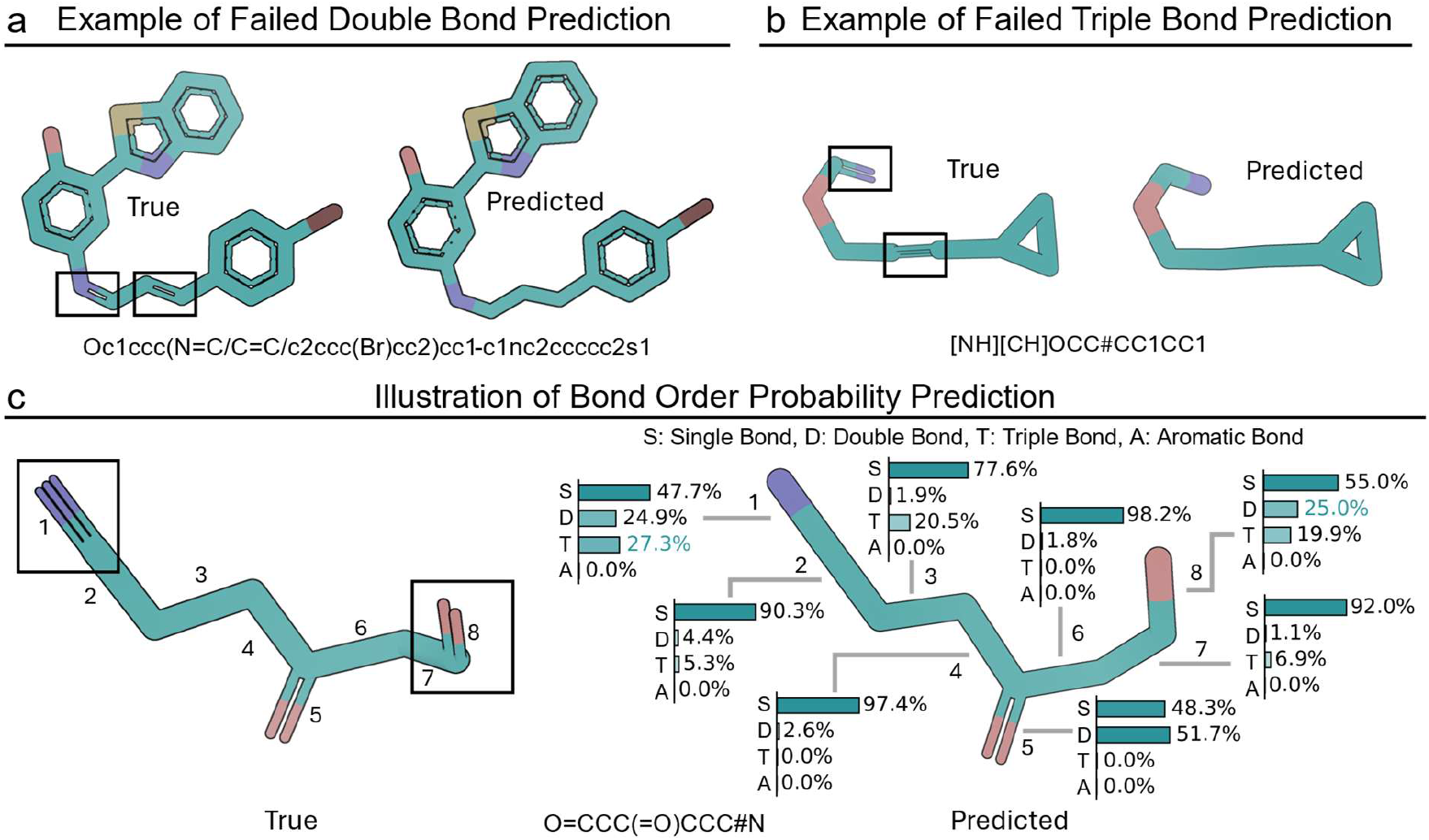
Illustration of Bond Order Probability Prediction. (a, b) Two examples of failed predictions for double and triple bonds. (c) Despite incorrectly predicting the triple and double bonds as single bonds, the probabilities for these bond orders remain non-negligible.

To illustrate this, we analyzed a representative compound with the SMILES O=CCC(=O)CCC#N, which contains two double bonds and one triple bond. We calculated the predicted probabilities for each bond in the molecule (Figure 5c). For example, bond 1, which is a true triple bond, is misclassified as a single bond. However, the model assigns a 27.3% probability to the triple bond class, compared to 47.7% for single and 24.9% for double. This relatively high probability for the correct class indicates that the model retains partial confidence in the correct prediction, even if it does not appear as the top choice. In contrast, bond 2, a correct single bond prediction, is associated with a confident 90.3% probability, showing the model’s strong certainty when the context is less ambiguous.

A similar pattern is seen in bond 8, which is a double bond but predicted as single. Here, the probability for double bond reaches 25.0%, while single and triple bonds receive 55.0% and 19.9%, respectively. These examples show that even when the final classification is incorrect, the predicted probability distribution often captures meaningful chemical ambiguity.

### Test on Kekulized Dataset

In previous scenarios, YuelBond was trained and evaluated using molecular representations that retained aromatic bonds as explicit aromatic types. However, in many cheminformatics applications, particularly in graph-based processing, it is common practice to kekulize aromatic systems, which is to convert aromatic bonds into alternating single and double bonds (Figure 6a). To assess the robustness of our model under such settings, we recompiled the GEOM dataset by kekulizing all aromatic bonds and retrained YuelBond using this modified dataset. We evaluated the performance across all three prediction scenarios. When reconstructing bonds from exact 3D coordinates, YuelBond achieved an average accuracy of 96.5%, precision of 96.5%, recall of 96.5%, and F1-score of 96.4% on the test set (Figure 6b; Table S4). In the scenario of CDG, performance remained high, with 91.9% accuracy, 91.0% precision, 91.9% recall, and an F1-score of 90.9% (Figure 6c; Table S5). In the most challenging bond order reassignment, YuelBond achieved 80.3% accuracy, 76.8% precision, 80.3% recall, and an F1-score of 76.6% (Figure 6d; Table S6).

**Figure 6.**
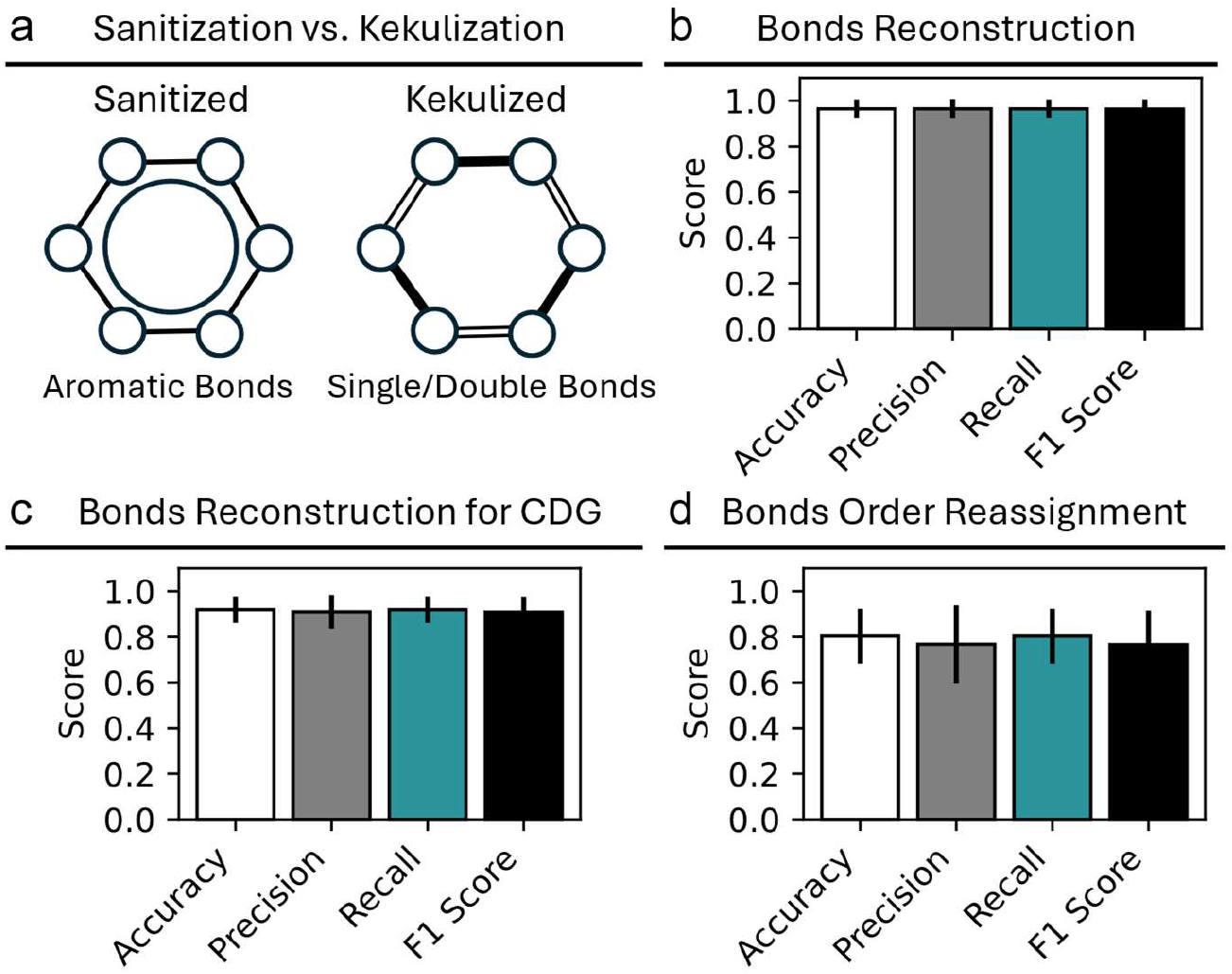
Performance of YuelBond on Kekulized Dataset. (a) In the Kekulized dataset, aromatic bonds are represented as alternating single and double bonds. (b) Performance of YuelBond on bond reconstruction from 3D coordinates. (c) Performance of YuelBond on bond reconstruction from CDG. (d) Performance of YuelBond on bond order reassignment.

Despite the structural transformation introduced by kekulization, the overall performance of YuelBond remains consistent with its original version trained on non-kekulized data, suggesting that YuelBond is not overly reliant on explicit aromatic bond annotations and can generalize across chemically equivalent but representationally different datasets. The minor drop in F1-score for bond order reassignment (from 78.3% to 76.6%) may reflect increased ambiguity in distinguishing alternating single and double bonds in kekulized rings. These results confirm that YuelBond maintains strong predictive performance even when trained on kekulized data.

## DISCUSSION

In the idealized case with accurate 3D atomic coordinates, YuelBond achieves near-perfect reconstruction. This reflects the high fidelity of 3D geometry in encoding bond-specific information such as distance and local spatial configuration. The slight conservatism in triple bond classification, manifested as high precision but lower recall, suggests a preference for avoiding false positives in ambiguous scenarios. It is a desirable behavior for downstream applications in molecular modeling, where incorrectly assigning a high bond order may introduce substantial artifacts.

The second set of experiments, which introduce geometric distortion to mimic the imperfections of CDG, underscores the resilience to structural noise. YuelBond not only remains functional under such perturbation but also maintains strong predictive power. While there is a drop in performance, especially for bonds that are sensitive to precise geometry such as double and triple bonds, the results remain significantly superior to RDKit. The RDKit benchmark shows that rule-based systems are fragile when assumptions about molecular structure are violated. YuelBond, trained on varied data and designed to incorporate learned spatial heuristics, proves more tolerant of noise, making it better suited for emerging applications in generative chemistry and structure prediction pipelines.

In the 2D-only setting, the problem shifts from one of geometric reasoning to one of chemical inference, as the model is forced to predict bond orders without direct access to atomic positions. Despite the lack of geometric cues, YuelBond performs still well, implying the capacity to learn structural priors purely from connectivity patterns. The disparity in performance across bond types, especially the lower recall for double and triple bonds, points to inherent ambiguities in 2D representations. Many different electronic or resonance states can yield the same connectivity graph, complicating prediction. The observed high accuracy but low recall for triple bonds in this scenario indicates a conservative bias of YuelBond, which tends to require strong evidence before assigning rare or chemically specific bond types.

The bond order probability highlights the ability of YuelBond to express prediction uncertainty. Unlike deterministic rule-based systems, YuelBond outputs a distribution over possible bond orders, which offers a more nuanced understanding of model confidence. This feature is especially valuable in ambiguous or edge cases, such as the analysis of a molecule with multiple functional groups, where assigning a single bond order may be chemically limiting. These probabilistic outputs can be incorporated into downstream workflows, e.g., during molecule generation, refinement, or validation, offering users a mechanism for prioritizing predictions, flagging uncertain assignments, or sampling alternative structures.

Taken together, YuelBond presents a unified framework capable of learning and generalizing chemical bonding rules across diverse input representations. It excels in accurate 3D settings, tolerates geometric distortion, adapts reasonably well to 2D graphs, and provides interpretable probabilistic predictions. Rather than aiming to replace rule-based methods entirely, YuelBond can be seen as a complementary tool, which brings the flexibility and contextual awareness of machine learning into scenarios where traditional approaches falter.

## METHODS

### Dataset and Preprocessing

We used the GEOM dataset for the training, validation and test. The dataset contains over 450,000 molecules. The GEOM dataset is a large-scale collection of molecular conformations annotated with energy and statistical weight information, designed to support machine learning applications in computational chemistry. It comprises approximately 37 million conformers for over 450,000 molecules, including 133,000 species from the QM9^43^ dataset and 317,000 species with experimental data related to biophysics, physiology, and physical chemistry. Each molecule’s conformers were generated using advanced sampling techniques and semi-empirical density functional theory (DFT) methods and further processed to explore conformational space. We used the conformer with the lowest energy of each molecule for the training, validation and test.

We extracted the SMILES strings and 3D conformer with the lowest energy of each molecule and stored them in an SQLite database for efficient retrieval. Each molecule undergoes two parallel processing streams: (1) sanitized (validated via RDKit’s standard chemical checks) and (2) Kekulized (aromatic bonds explicitly represented as alternating single/double bonds). For each molecule, atomic positions are extracted from the conformer, while atoms and bonds are encoded as one-hot vectors using predefined mappings. Invalid and large molecules (e.g., those with <2 atoms or >150 atoms) are filtered out.

Molecular graphs are constructed by representing atoms as nodes (with one-hot features and 3D coordinates) and bonds as edges (with one-hot bond types). Key molecular features, including atomic positions, one-hot encoded atom types, and bond types, are extracted. In the scenario of predicting the bond order of the molecule from 3D coordinates, edges are created between atoms within a 3 Å distance cutoff. In the scenario of predicting the bond order of the molecule from the connectivity of the atoms, edges are created as the bonds in the molecule, but the bond order is not included. In the scenario of predicting the bond order of CDG, Gaussian noise (σ = 0.2 Å) is added to the atomic positions, and then edges are created between atoms within a 3 Å distance cutoff.

Finally, variable-sized molecules are padded to the largest molecule in each batch, with node masks and edge masks marking valid nodes/edges. Processed datasets are cached in files for efficient reloading.

### Architecture of the Bonds Reconstruction Model

The neural network is a graph neural network designed to predict bond orders in molecular compounds by processing graph-structured data, where nodes represent atoms and edges represent bonds. The model begins by embedding input node and edge features into a higher-dimensional space using linear transformations. The core of the network is a stack of message passing layers, which iteratively refine node and edge features through neighborhood aggregation. Each message passing layer contains two key components: an edge model and a node model. The edge model processes edge features by combining source and target node features with existing edge attributes, passing them through a multi-layer perceptron (MLP)^41^ with layer normalization and a SiLU^42^ activation function. An innovation is the incorporation of existing edge features into the edge model. Previous GNNs only use the features of the connected nodes to update the edge features. In this work, we use the concatenation of the features of the connected nodes, the distance between the nodes, and the previous edge features to update the edge features. This way, the model can expand the receptive field more effectively. The node model aggregates incoming edge features for each node, combines them with the node’s current features, and applies another MLP to update the node representation. The forward pass handles batched graphs by merging them into a single large graph, applying masked operations to respect variable-sized inputs, and then splitting the results back into batches. The final output is produced by projecting the hidden features to the desired output dimensions (e.g., bond orders for edges).

### Implementation of Graph Neural Network with Edge Updates

We model the molecular system as a fully connected undirected graph *G* = (*V, E*), where each node *v*_*i*_ ∈ *V* represents an entity (e.g., an atom or residue) with a feature vector *h*_*i*_ ∈ ℝ^d^, and each edge *e*_*ij*_ ∈ *E* represents an interaction or relationship between nodes *v*_*i*_ and *v*_*j*_, with associated edge attributes *e*_*ij*_ ∈ ℝ^k^.

Our framework builds on the message-passing neural network (MPNN)^40^ paradigm but introduces a dedicated edge update mechanism that plays a central role in learning the relational structure of the data. Unlike standard GNNs that update edge features implicitly or keep them static, we explicitly model and update *e*_*ij*_ during each layer of the network.

At each layer *l*, node features and edge features are updated as follows:

Message construction: For each pair of nodes (*i, j*), a message is constructed using a learned function ϕ_m_:

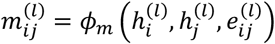

Edge update: The edge features are updated using the current node representations and previous edge features through a function ϕ_e_:

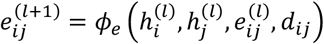

Node update: Each node aggregates incoming messages from its neighbors 𝒩(*i*) using an aggregation function ρ, followed by an update function ϕ_h_:

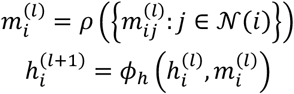

All functions *ϕ*_*m*_, *ϕ*_*e*_, and *ϕ*_*h*_ are implemented as two-layer (MLPs with SiLU^42^ activations and layer normalization. To ensure stability during training, we use residual connections where applicable.

### Training Process

The model is trained using PyTorch^44^ Lightning, which streamlines the training loop, validation, and logging. The training step involves passing molecular graphs through the GNN to predict bond orders, followed by computing the loss between predicted and true bond types. The loss function is masked cross-entropy, where only valid edges (non-padded bonds) contribute to the gradient updates. The optimizer used is AdamW^45^ with a learning rate of 10^−4^, AMSGrad enabled, and weight decay (10^−12^) for regularization. Training metrics (e.g., loss) are logged at specified intervals and tracked using Weights & Biases^46^ for visualization. The model supports data augmentation and processes batches of variable-sized graphs using a custom collate function to handle padding and masking.

### Loss Function

The objective of the model is to accurately predict the presence and type of bonds (or edges) between node pairs in a graph. To this end, we define a supervised learning loss based on the categorical cross-entropy between the predicted bond type distribution and the ground-truth bond type labels.

Let *p*^*ij*^ ∈ ℝ^*c*^ be the predicted probability distribution over *C* bond types for the edge between nodes *i* and *j*, and let *y*_*i*_ ∈ {0,1}^*c*^ be the one-hot encoded ground truth label. The edge prediction loss is given by:

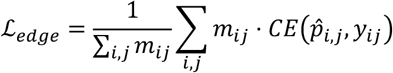

where 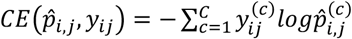 is the standard cross-entropy loss for a single edge. *m*_*ij*_ ∈ {0,1} is a binary mask indicating whether the edge (*i, j*) is valid (1) or padded/invalid (0). The denominator ∑_*i,j*_ *m*_*ij*_ normalizes the loss by the number of valid edges to ensure stability regardless of graph size or padding.

The masked and normalized formulation prevents the model from being biased by graph sparsity or the inclusion of padded entries during batching. It also ensures consistent gradients during training across variable-sized graphs.

### Evaluation Metrics

The model’s performance is evaluated using standard multi-class classification metrics: accuracy, precision, recall, and F1-score. They are computed on a per-molecule basis and aggregated across the test set. The classification targets are the bond types between atom pairs, such as single, double, triple, and aromatic. Padded or invalid edges are masked and excluded from all metric computations.

Let ŷ_*ij*_ ∈ (1, …, *C*) denote the predicted bond type for atom pair (*i, j*), *y*_*ij*_ the corresponding ground-truth bond type, *m*_*ij*_ ∈ {0,1} the mask for valid edges, *C* the number of bond classes, 𝕀 the indicator function, then:

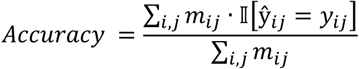

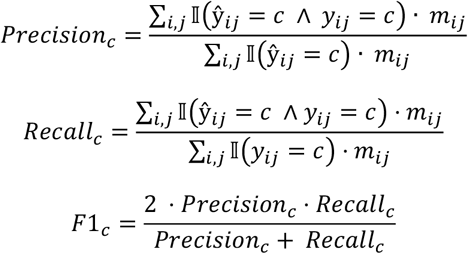

### Baseline Method: RDKit’s DetermineBonds Algorithm

To assess the performance of YuelBond, we benchmark it against the built-in bond type assignment function of RDKit^33^, *rdkit*.*Chem*.*rdDetermineBonds*.*DetermineBonds*. This method is a widely-used rule-based algorithm that infers bond orders directly from molecular 3D geometries. Internally, it leverages the xyz2mol^34^ program, which applies a series of chemically motivated heuristics to reconstruct the bond network of a molecule from atomic coordinates.

The xyz2mol algorithm proceeds in multiple stages. First, it detects bonded atom pairs based on interatomic distances and covalent radii thresholds. Next, it computes potential valence states for each atom and iteratively searches for a valid bond order configuration. This process prioritizes chemical plausibility by enforcing standard valency constraints and ensures that no atom exceeds its typical bonding capacity. For cases involving multiple plausible configurations, particularly in conjugated or aromatic systems, xyz2mol employs additional reasoning strategies, including Hückel theory or graph-theoretic techniques, to resolve ambiguities and assign aromaticity. While *DetermineBonds* offers a fast and interpretable solution, its performance is inherently limited by the rigidity of handcrafted rules and its reliance on idealized geometry.

## ACKNOWLEDGMENTS

We acknowledge support from the National Institutes for Health 1R35 GM134864, the Huck Institutes of the Life Sciences, and the Passan Foundation. The content is solely the responsibility of the authors and does not necessarily represent the official views of the NIH. This project was supported by the Penn State College of Medicine’s Artificial Intelligence and Biomedical Informatics Program.

## DATA AND SOFTWARE AVAILABILITY

Source codes and test data are deposited at: https://bitbucket.org/dokhlab/yuel_bond.

## SUPPORTING INFORMATION AVAILABLE

The supporting information provides Table S1-S6.

## DECLARATION OF INTERESTS

The authors declare no competing financial interest.

## AUTHOR CONTRIBUTIONS

Jian Wang contributed to the conceptualization, methodology, model development, data analysis, writing of the original draft, and visualization of the study. Nikolay V. Dokholyan provided supervision, resources, writing review and editing, project administration, and funding acquisition.

## Notes

https://bitbucket.org/dokhlab/yuel_bond

### Competing Interest Statement

The authors have declared no competing interest.

