## Supplemental Information for "Multimodal Bonds Reconstruction Towards Generative Molecular Design"

**Table S1. Test Performance of Reconstructing Bonds from Accurate 3D Coordinates.**

| | Accuracy | | Precision | | Recall | | F1 Score | |
|---|---|---|---|---|---|---|---|---|
| | Average | Std. Dev. | Average | Std. Dev. | Average | Std. Dev. | Average | Std. Dev. |
| All | 0.983 | 0.034 | 0.988 | 0.025 | 0.983 | 0.034 | 0.984 | 0.031 |
| Single | 0.984 | 0.030 | 0.954 | 0.110 | 0.956 | 0.102 | 0.951 | 0.102 |
| Double | 0.996 | 0.012 | 0.954 | 0.168 | 0.926 | 0.207 | 0.914 | 0.221 |
| Triple | 0.999 | 0.007 | 0.988 | 0.078 | 0.894 | 0.294 | 0.894 | 0.292 |
| Aromatic | 0.986 | 0.030 | 0.902 | 0.251 | 0.955 | 0.115 | 0.894 | 0.250 |

**Table S2. Test Performance of Reconstructing Bonds from CDG.**

| | Accuracy | | Precision | | Recall | | F1 Score | |
|---|---|---|---|---|---|---|---|---|
| | Average | Std. Dev. | Average | Std. Dev. | Average | Std. Dev. | Average | Std. Dev. |
| All | 0.927 | 0.082 | 0.936 | 0.075 | 0.927 | 0.082 | 0.927 | 0.081 |
| Single | 0.935 | 0.074 | 0.834 | 0.210 | 0.860 | 0.183 | 0.833 | 0.189 |
| Double | 0.983 | 0.024 | 0.859 | 0.303 | 0.560 | 0.434 | 0.563 | 0.431 |
| Triple | 0.998 | 0.013 | 0.865 | 0.338 | 0.640 | 0.466 | 0.599 | 0.479 |
| Aromatic | 0.952 | 0.066 | 0.780 | 0.316 | 0.832 | 0.230 | 0.755 | 0.316 |
| RDKit | 0.651 | 0.244 | 0.700 | 0.302 | 0.651 | 0.244 | 0.641 | 0.277 |

**Table S3. Test Performance of Reassigning Bond Orders for 2D Atom Graphs.**

| | Accuracy | | Precision | | Recall | | F1 Score | |
|---|---|---|---|---|---|---|---|---|
| | Average | Std. Dev. | Average | Std. Dev. | Average | Std. Dev. | Average | Std. Dev. |
| All | 0.801 | 0.199 | 0.811 | 0.216 | 0.801 | 0.199 | 0.783 | 0.219 |
| Single | 0.804 | 0.197 | 0.789 | 0.223 | 0.917 | 0.155 | 0.824 | 0.179 |
| Double | 0.946 | 0.066 | 0.836 | 0.317 | 0.493 | 0.446 | 0.494 | 0.440 |
| Triple | 0.992 | 0.034 | 0.750 | 0.407 | 0.296 | 0.450 | 0.267 | 0.429 |
| Aromatic | 0.861 | 0.187 | 0.853 | 0.289 | 0.731 | 0.334 | 0.716 | 0.342 |

**Table S4. Test Performance of Reconstructing Bonds from Accurate 3D Coordinates in the Kekulized Dataset.**

| | Accuracy | | Precision | | Recall | | F1 Score | |
|---|---|---|---|---|---|---|---|---|
| | Average | Std. Dev. | Average | Std. Dev. | Average | Std. Dev. | Average | Std. Dev. |
| All | 0.965 | 0.041 | 0.965 | 0.043 | 0.965 | 0.041 | 0.964 | 0.043 |
| Single | 0.966 | 0.041 | 0.918 | 0.097 | 0.944 | 0.082 | 0.930 | 0.086 |
| Double | 0.965 | 0.041 | 0.831 | 0.225 | 0.773 | 0.242 | 0.786 | 0.238 |

| | | | | | | | | |
|---|---|---|---|---|---|---|---|---|
| Triple | 0.999 | 0.008 | 1.000 | 0.000 | 0.887 | 0.294 | 0.897 | 0.283 |

**Table S5. Test Performance of Reconstructing Bonds from CDG in the Kekulized Dataset.**

| | Accuracy | | Precision | | Recall | | F1 Score | |
|---|---|---|---|---|---|---|---|---|
| | Average | Std. Dev. | Average | Std. Dev. | Average | Std. Dev. | Average | Std. Dev. |
| All | 0.920 | 0.058 | 0.910 | 0.074 | 0.920 | 0.058 | 0.909 | 0.068 |
| Single | 0.921 | 0.058 | 0.822 | 0.129 | 0.922 | 0.087 | 0.865 | 0.099 |
| Double | 0.935 | 0.049 | 0.733 | 0.313 | 0.430 | 0.341 | 0.474 | 0.347 |
| Triple | 0.998 | 0.014 | 0.818 | 0.375 | 0.644 | 0.472 | 0.570 | 0.484 |

**Table S6. Test Performance of Reassigning Bond Orders for 2D Atom Graphs in the Kekulized Dataset.**

| | Accuracy | | Precision | | Recall | | F1 Score | |
|---|---|---|---|---|---|---|---|---|
| | Average | Std. Dev. | Average | Std. Dev. | Average | Std. Dev. | Average | Std. Dev. |
| All | 0.803 | 0.120 | 0.768 | 0.171 | 0.803 | 0.120 | 0.766 | 0.150 |
| Single | 0.803 | 0.120 | 0.821 | 0.115 | 0.946 | 0.078 | 0.874 | 0.080 |
| Double | 0.815 | 0.121 | 0.716 | 0.309 | 0.385 | 0.333 | 0.424 | 0.334 |
| Triple | 0.989 | 0.041 | 0.783 | 0.413 | 0.144 | 0.334 | 0.145 | 0.337 |
